# Evolution of primate protomusicality via locomotion

**DOI:** 10.1101/2020.12.29.424766

**Authors:** David M. Schruth, Christopher N. Templeton, Darryl J. Holman, Eric A. Smith

## Abstract

Animals communicate acoustically to report location and identity to conspecifics. More complex patterning of calls can also function as displays to potential mates and as territorial advertisement. Music and song are terms often reserved only for humans and birds, but elements of both forms of acoustic display are also found in non-human primates. While theories on proximate functions abound, ultimate drivers of specific call structures are less well understood. We hypothesized that spatio-temporal precision in landing during perilous arboreal locomotion favored the evolution of musical calling in early primates—vastly preceding the origin of more music-like behavior in hominoids and subsequent emergence of music in later hominids. We test this locomotion-based hypothesis on the origins of proto-musicality using spectrographic depictions of vocal repertoires of modern day primates and corresponding estimates of locomotor activity. Phylogenetically controlled regression analysis of 54 primate species reveals that arboreal locomotion and monogamy are robust influences on complex calling patterns while controlling for other socioecological variables. Given that these findings rest primarily upon a handful of deep branching points in the primate tree, we conclude that this coevolution likely occurred very slowly, occupying on the order of tens of millions of years.

**License:** Attribution-NoDerivatives 4.0 International (CC BY-ND 4.0)

## Introduction

The origins of human music are confounded by a lack of consensus on theoretical evolutionary mechanisms and a seemingly unavoidable circularity in definitions (Schruth, Templeton and Holman, 2019). Humans are complex musical beings with an unusual ability to adapt in cultural as well as genetic, cognitive, and ecological ways (Smith, 2011). Many correspondingly plausible adaptive mechanisms have been proposed including: sexual [or mate] choice (Darwin, 1871; Miller, 2000), credible signaling (Mehr *et al*., 2020), coalitional or group selection (Hagen and Bryant, 2003), cultural evolution (Savage, 2019), gene-culture co-evolution (Cross, 2003), and epigenetic modification (Mehr *et al*., 2020). Similarly unresolved are reasonable, albeit western (Jacoby *et al*., 2020), definitions of the musical units of investigation including: song as relatively complex calls used in conspecific interactions (Beecher and Brenowitz, 2005), complex acoustic display (Templeton *et al*., 2011), or learned complex calls (Fitch, 2015); music as information rich holistic patterns (Roederer, 1984), or creative orderly, organized, structured sequences with repeatable distinctive patterns (Marler, 2000); and musicality as a neurobiologically constrained and spontaneous capacity to receive and produce such stimuli (Morley, 2002, 2012; Honing *et al*., 2015). A lack of clarity concerning the whats (outcomes and inputs) and hows (level, unit, tempo, and mode) of the evolution of musicality, however, has thus far stifled rigorous testing of origins theories.

Similar to ambient noise masking of aquatic signals (Balebail and Sisneros, 2020), vegetative obstruction is thought to ecologically select for salient calls in arboreal animals (Morton, 1975; Krause, 1993; Slater, 2000). Yet human musicality presents a puzzle as we do not typically face similar constraints of arboreality, having adapted to more open habitats since the middle Pleistocene (Grove, 2011). While there are many (mostly) arboreal species that exhibit music-like behavior, humans are singular in being habitually terrestrial (Brown and Jordania, 2013). Animals are known to use calls which contain song-like structures to localize themselves to conspecifics (Pollock, 1986; Catchpole and Slater, 1995). They have further compulsion towards more exceptional vocal displays—ranging from asserting unique identity to specializing features of their territorial advertisements (Goustard, 1984; Pollock, 1986; Cooney and Cockburn, 1995). In the light of ecological resource instability (Mattison *et al*., 2016) the case for musicality as a territorial signal in the most recent, hominin, environment of evolutionary adaptedness is debatable.

Plausible theories on music origins in humans range from infant attention (Trehub and Trainor, 1998; Dissanayake, 2000) to group communication (Brown, 2000; Hagen and Bryant, 2003). Darwin suggested that musical notes and rhythm functioned as part of courtship (Darwin, 1871), a theory others have endorsed (Miller, 2000; Dunbar, 2012). Until recently, the definition of music itself has been confounded with context, such as culture, materials, and group setting (Nettl, 2000), rendering independent efforts to understand functional origins challenging (Schruth, Templeton and Holman, under review). And while it has been quite common to use the term *function* in a way that is nearly synonymous with *proximate context* (Pollock, 1986; Cooney and Cockburn, 1995; Templeton *et al*., 2011; Mehr *et al*., 2018), research into ultimate evolutionary influences is rare. We suggest that an investigation into these ultimate adaptive causes of hominin musicality could benefit from ecological and signaling theory insights on primate behavior whereby contexts are understood separately from the “acoustic features themselves” (Merriam and Merriam, 1964). To begin addressing the possible ecological drivers of a pre-hominin musicality, we examine vocalizations of extant primates and their possibly functional relationships with discontiguous locomotion through arboreal substrate. Specifically, we hypothesized that the bifurcating topologies of primates’ arboreal habitats may not only have selected for the cognition necessary for *survival* in such precarious settings (Collins, 1921; Clark, 1959), but also that they may have favored the development of signals as indicators of these underlying abilities to conspecifics, both to potential mates and resource competitors.

We leverage the overarching theoretical framework of behavioral ecology to model the fit of (e.g. musical) behavior to (e.g. an arboreal) environment—assuming a process of natural selection by both physical surroundings and the behavior of other organisms (Fox and Westneat, 2010). Additionally, we leverage the concept of mate choice—the full cycle including courtship, copulation, fertilization, and parenting all recently acknowledged to represent a behavioral continuum (Dissanayake, 2008; Brooks *et al*., 2010; Savage, 2019)—to help in absolving misunderstandings regarding which social factors are most important in rewarding proto-musical behavior. We know, for example, that social monogamy has a strong association with musical behavior (Haimoff, 1986) but such proximate connections are often conflated with ultimate causality (Mehr *et al*., 2020). Accordingly, we focus on natural selection for robust survival traits (e.g, locomotion) that are signaled by *senders* (of musical displays), rather than more proximately subvertable selection (e.g. via sexual choice) by *receivers*, although both are crucial components.

We build on hypotheses that musical displays could demonstrate full maturation of generalized [dimensional] comparison abilities (Roederer, 1984) and [vocal-fold] motor control (Calvin, 1982; Roederer, 1982; Pinker, 1997)—capabilities useful for [visual focus and other] fine-motor tasks (Sacks, 2007). Beyond these proposed sensory-motor links, it is also possible that many auditory-musical spectrum behaviors are associated with spatial cognition (Dehaene *et al*., 2003; Harris and Miniussi, 2003; Farrell *et al*., 2012) such as auditory interval with verticality perception (Melara and O’Brien, 1987; Rusconi *et al*., 2006; Bonetti and Costa, 2019). For the proto-musical calling of primates, we are most interested in correlates of melodic processing. Brain imaging studies typically locate music and melody perception in higher-cortical areas such as the temporal gyrus (Morley, 2002, 2012), but the limbic system has also recently been implicated (Harvey, 2017). These mid and hind brain areas, including the hippocampus, basal ganglia, and cerebellum, are thought to participate in melodic binding (Fernández, 2015). The hippocampus, in turn, also serves as a key facilitator of spatial cognition (Save and Poucet, 2000). Similar connections between song and equivalent brain structures in birds have also recently been observed (Nicholson, Roberts and Sober, 2018; Pidoux *et al*., 2018). It is possible that these underlying spatial proficiencies, and corresponding spatio-sensory motor control abilities, could have been evolutionary selected in the *sender* to indirectly signal such qualities to conspecific receivers of musical calls. Senders and receivers could mutually benefit from the honesty of such signals via resource spacing, conflict avoidance, and mating potential. Dimensional precision for difficult aerial sensory-motor tasks (e.g. landing with velocity in complex canopy habitats composed of tenuous branches) could efficiently be signaled to others (likely including kin) within a breeding deme. This mode of signaling avoids venturing onto the forest floor or using diffused chemical, visually occluded, or otherwise ineffectual signals (Slater, 2000). In summary, we propose that arboreal primates, intent on avoiding terrestrial predation (Schruth and Jordania, 2020), frequently became at least moderately airborne in order to traverse gaps in substrate—and that the selection for corresponding (e.g. ocular) motor control and spatial cognition (e.g. resolving arbitrary branch shapes) for landing such bouts, maintained the honesty of such precise vocal signals.

The evidence for musical behavior in the archaeological record is slim (D’Errico *et al*., 1998) and virtually non-existent in the paleontological record, making the testing of adaptive origins theories intractable. Alternatively, researchers might utilize modern day analogs to either reconstruct or statistically infer what ancestral calls may have been like (Wich and Nunn, 2002). Unfortunately, only a handful of primate species are considered “musical” (Geissmann, 2000) and such binary assessments make ancestral reconstruction statistically insoluble. In addition to traditional binary classifications, we used a continuous measure of musicality, the acoustic reappearance diversity index [ARDI] (Schruth, Templeton and Holman, 2019). ARDI is an estimate of the number of reappearing syllables within a call type (a rough proxy for protomusical behavior) and was derived from analysis of ethnomusicalogically prevalent acoustic features observed in primate calls (Schruth, Templeton and Holman, under review). We investigate this theory by analyzing non-human primate data within the evolutionary testing framework of phylogenetically controlled regression modeling.

## Materials and Methods

We collected spectrographic vocal repertoires from the literature by searching Web of Science Citation Index (Garfield, 1970) using the partial search terms “spectro* AND primate*AND <genus>“ with asterisks indicating wild cards. Subsequent searches via google scholar (Acharya and Verstak, 2004) helped to fill in gaps by finding studies on species from genera with sparse representation in the larger dataset. In total 832 vocalizations from 60 species were collected corresponding to 39 genera and all but one primate family. Spectrograms were cropped out of their axes, renamed, and anonymized before scoring.

Scoring took place over the course of two days using bird call examples as training materials. Each of the five scorers had a different ordered spreadsheet of calls and scored, on a 1-10 scale, six different acoustic parameters: tone, interval, rhythm, repetition, transposition, and syllable count. Details of this scoring protocol are available online (Schruth and Holman, 2020). Scores were reliable across scorers with values ranging from 0.7 to 0.9 using Cronbach’s alpha measure (Cronbach, 1970). These scores were then converted to a single number per vocalization via averaging between the scorers resulting in a total of 832 scores for six different parameters. This matrix was then input into PCA software (R Core Team, 2018) to help reduce the six variables into a more manageable number of variables for further analysis. PCA results suggested retaining (Jolliffe, 1972) repetition, transposition, and syllable count—the last of which is a commonly measured feature of avian songs (Wildenthal, 1965; Botero *et al*., 2008). We reasoned that repetition and transposition are mutually exclusive and could be combined into a single measure of *redundancy*. Reappearance, in turn, was then multiplied by the unique syllable count to create a reappearance weighted measure of spectral shape diversity. This acoustic reappearance diversity index [ARDI] corresponded well to vocalizations designated by primary researchers as being “song” or “musical.” Since rhythm was not retained by our PCA reduction procedure, however, the resulting index is admittedly better at capturing more transpositionally melodic calls over those that are more rhythmically complex (Schruth, Templeton and Holman, under review). Details are available in another manuscript (Schruth, Templeton and Holman, 2019) but data and spectrograms are available online (Schruth, 2019).

Locomotion data was collated from the primate literature in a search procedure analogous to that employed for the spectrographic data—using “locomot*primate*<genus>“ search terms—as detailed above. In total the locomotion data set contained 54 different genera and 112 species. Studies were required at a minimum to have a quantitative estimate for leaping. But all other modes of locomotion were tabulated as well. Leaping and swinging percentages were cross-checked and verified against secondary compilations of locomotion (Rowe and Meyers, 2017). Leaping was coded as a composite variable combined with jump, air, and drop modes. Swinging was also composite with armswing and other suspensory modes.

We used regression (R Core Team, 2018) to compare our ARDI proto-musicality variable with a handful of candidate socioecological and locomotion variables. We calculated independent contrasts (Felsenstein, 1985) on each of these variables so as to control for non-independence of data collected at terminal nodes of the primate evolutionary tree, as closely related species shouldn’t be considered independent points (Felsenstein, 1985). These regression results were further compared with PGLM (*caper* v. 0.5.2) regression (Orme *et al*., 2013) on the same data using the same tree. We permuted over all possible modeling variable combinations—of *wooded, group size, monogamy*, as well as *leaping* and *swinging—*and averaged the resulting maximum likelihood estimates to obtain a static set of tree transformation parameters (kappa=2.5, lambda=0.2, delta=1.3) for the final PGLM analysis.

## Results

Our results suggest that aerially discontiguous forms of locomotion, such as leaping and swinging, as well as social monogamy are each credibly associated with musical calling, but are somewhat contingent upon the specific method of phylogenetic control employed. Monogamy and locomotion contrasts exhibited the largest positive associations with protomusical calling as assessed by ARDI (Table 1, Figs 1 & 2). Monogamous species averaged nearly an entire additional reappearing syllable compared to non-monogamous species (β∼0.9; p<0.03). Leaping and swinging had nearly two fold greater effects than monogamy (for IC and PGLM respectively)—with an additional reappearing syllable in the most song-like call for every half range increase in leap bouts (IC; β∼2.0; p<0.05) and swing bouts (PGLM; β∼1.8; p<0.02). Further evidence of the importance of the monogamy and locomotion variables is seen in the fact that they were both significant under all models reported (Table 1) including the model with the highest R^2^ and that with the lowest AIC (Table 2), although only simultaneously for both methods in the locomotion only model. Wooded habitat and group size had positive associations but were not significantly different from null. The locomotion and mating model with a relatively high explanation of variance (26% and 38%) and amongst the lowest AIC (155 and 138), respectively, is the most informative model for the purposes of this study. These results were even more striking, however, when the two locomotion measures were added together (PGLM; β∼1.5; p<0.03), while using a binary “musical” outcome variable (PGLM, p<0.01, for swing; IC, p<0.02, for leap), or under index compositions that included an even greater number of musical features, such as those incorporating both rhythm and tone. Thus our findings of a relationship between locomotion and proto-musicality, using ARDI, are much more conservative by only including the features of transposition, repetition, and syllable count.

**Table 1.**
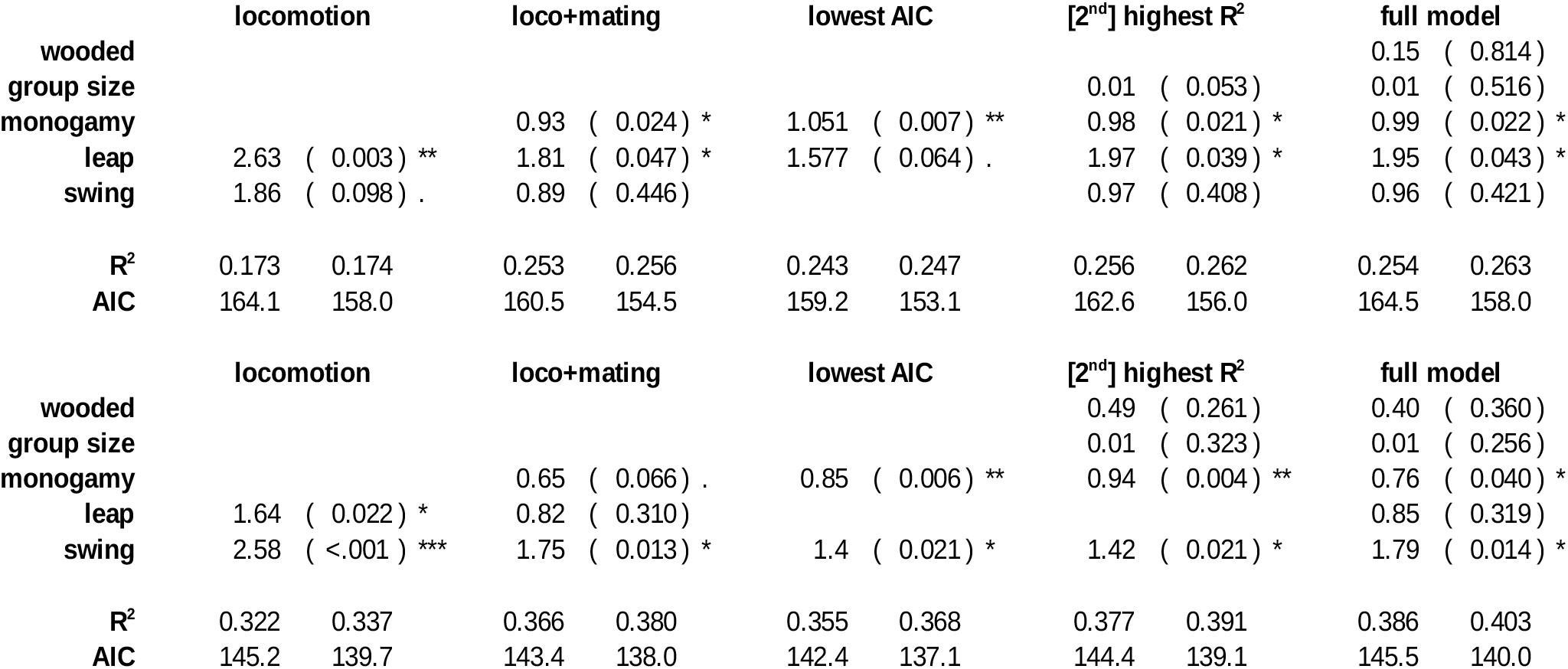
Multiple regression results for the contrasts between ARDI and various predictor. This table of results includes multivariate regressions, “full model” (right) and all others (left), and reflects modeling based on the coefficient of determination (R^2^) and Akaike’s information criterion (AIC). The top and lower table correspond to independent contrasts and PGLM (using kappa=2.5, lambda=0.21, delta=1.33) regression methods respectively. P-values are contained within parenthesis with adjacent stars and periods indicating levels of significance (^**^=0.01, *=0.05, and .=0.1). The greater significance of leaping under PGLM and swinging under IC, likely stems from differences in how the underlying tree is allowed to transform and adjust (e.g. the ML optimized kappa, lambda, and delta) in compensating for the relative rarity of swinging primates.

**Fig 1.**
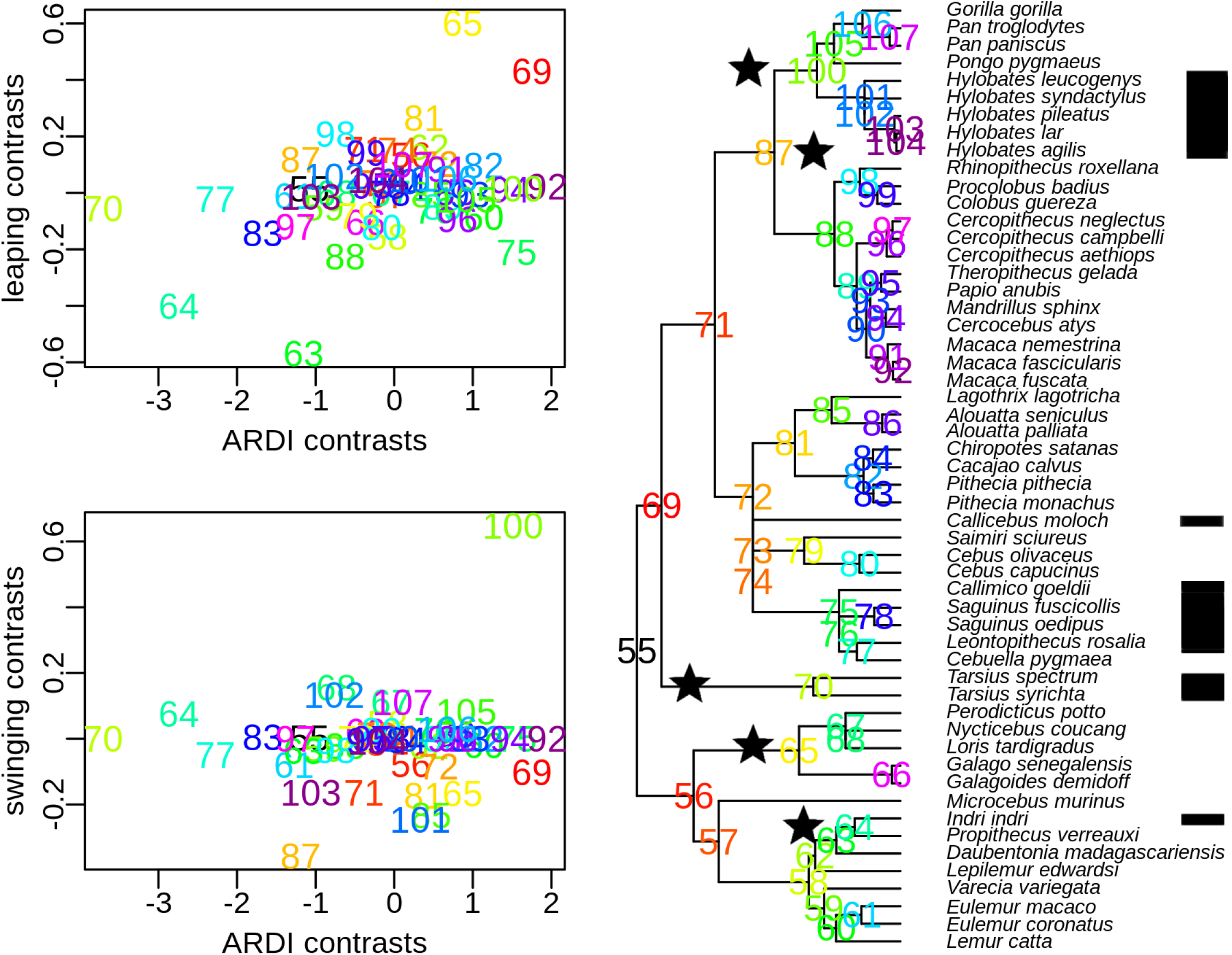
Plots of independent contrasts—between acoustic reappearance diversity [ARDI] and two locomotion predictors—and the corresponding phylogenetic tree. Only a handful of ancient branching points in the primate tree drive these two significant correlations. Four of these divergences are between leaping species (stars at bottom) and two are from brachiating species (stars at top). With the exception of *Platyrrhines*, all of these key divergences match up quite well with species thought to be musical (black bands on right) by previous investigators (Geissmann, 2000). Contrast 69 is the deepest branching point (at ∼60 MYA) and happens to represent the main split between *Tarsiiformes* and *Anthropoids*. Contrasts 87 and 100 are also rather old (∼30 and ∼20MYA), representing the split between *Cercopithecoids* and *Hominoids*, and *Hylobatids* from *Hominids*. Contrast 100 represents the significant difference between the *Hylobatids* (arboreal brachiators) and *Hominoids* (partially terrestrial knuckle-walkers). The rest of the main significance-driving branching points (65, 64, and 63) all relate to splitting *Indri* and *Galagoidae* off from *Pottos*. Note formal phylogenetic names are used in this caption while common names are used in the discussion section of the main text. Numeric labels of the internal nodes of the phylogenetic tree start just after the 54 extant primates (at the tree tips). Contrasts are calculated as differences in raw values between values at descendant branches: *max*(*ARDI*)_*SP1*_ *- max*(*ARDI*)_*SP2*_.

**Fig 2.**
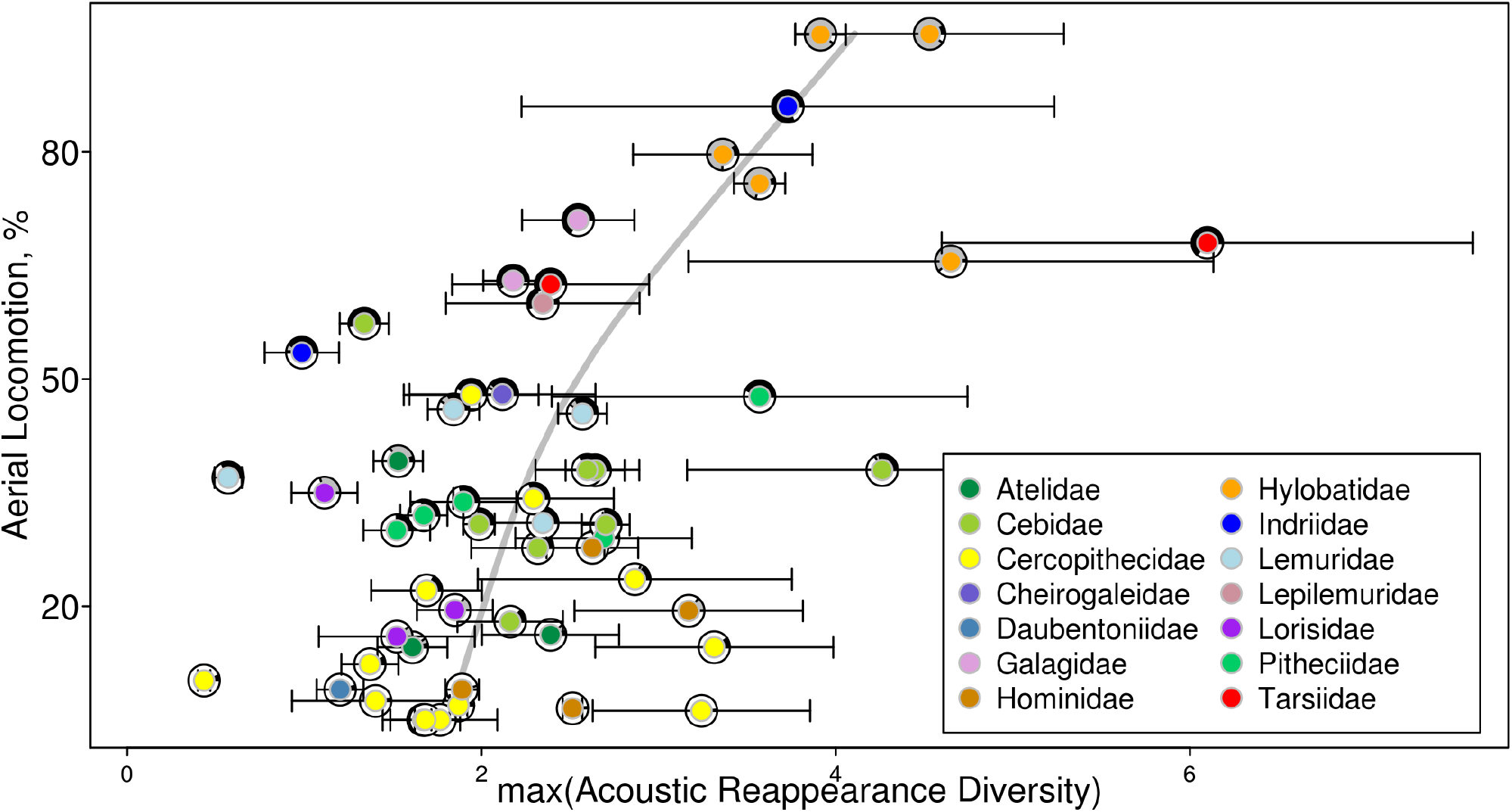
A scatterplot of reappearance diversity versus precision landing locomotion forms. Precision limb landed forms of locomotion leaping and swinging are added together to comprise the total aerial-spectrum locomotion percentage and are plotted against max (±SE) reappearance diversity [ARDI] scores on primate spectrograms for each species (*n*=54). The standard error for each reappearance diversity score was estimated via bootstrap by taking the standard deviation of the max estimates for 10,000 different samplings (with replacement) of all vocalization-level reappearance diversity scores for each species. A smooth spline (gray line) was fit to the data (using 3 degrees of freedom). Point colors indicate taxonomic family membership as specified by the key. Pie chart rings around each point represent the swing and leap percentages as grey and black.

**Table 2.**
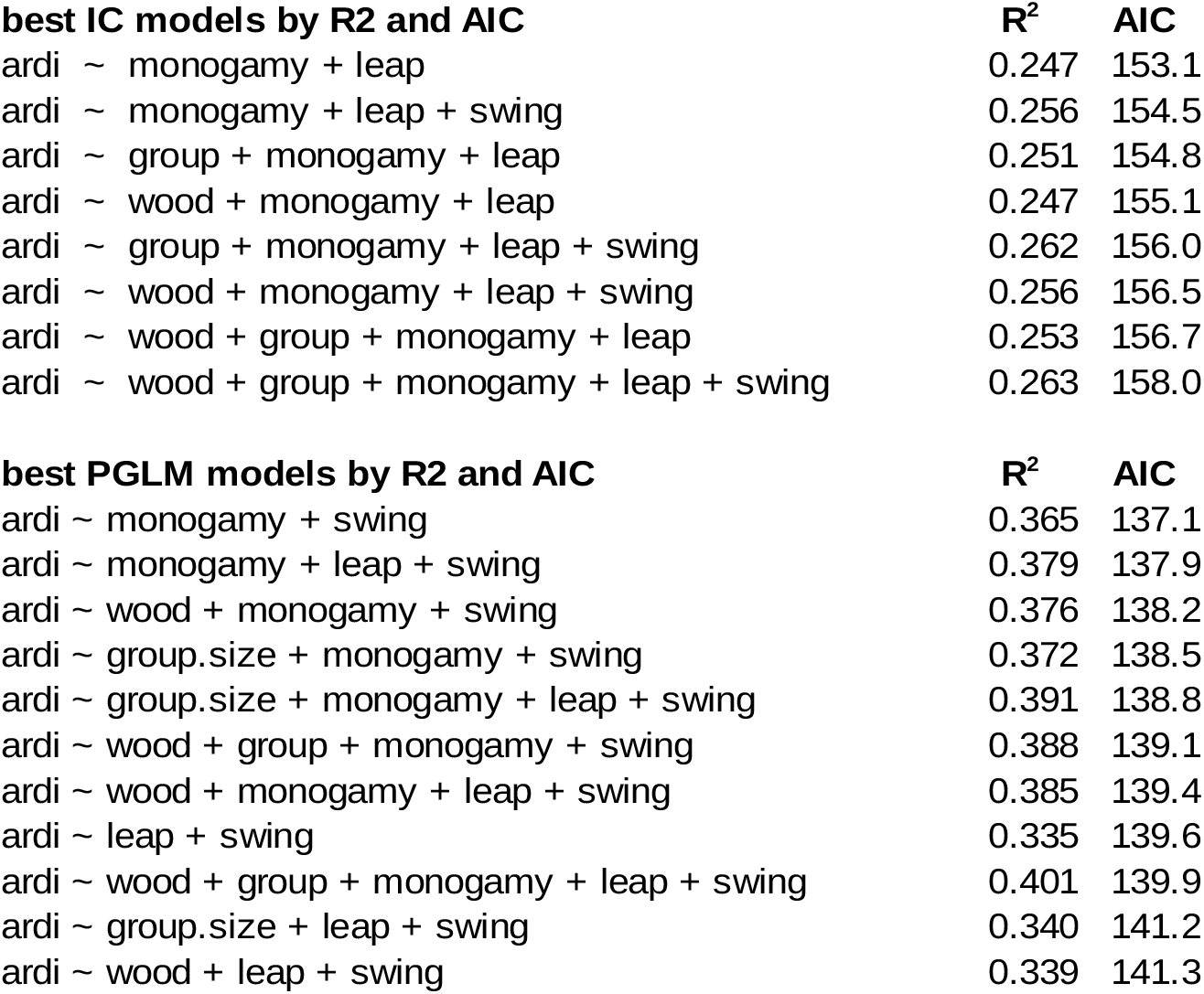
A list of tested statistical models and their corresponding AIC and R^2^ values. Models were filtered by those with R above 20% explained variance and sorted by increasing AIC.

## Discussion

The primary conclusion of our study—that arboreal pressures on primates may have driven the co-evolution of aerial spectrum locomotion (e.g. leaping and swinging) with song-like, proto-musical calling (Fig 2)—is largely derived from a handful of remarkable contrasts in each of these behaviors between phylogenetic neighbors (Fig 1). Specifically, the highly musical and frequently leaping Tarsiers and Indri (Fig 1: contrasts 72 and 67) and the quiet and non-leaping Loris and Aye-Aye (Fig 1: contrasts 68 and 66) constitute the four main drivers of the positive regression line trend in the leaping contrasts plot. More surprisingly, Galagos opposite Lorises (Fig 1: contrast 68) and *Pitheciidae*, such as titis, sakis, and uakaris, (opposite *Atelidae*, such as howler, spider, and woolly monkeys) emerge as relatively musical species as well (Fig 1: contrast 84).

The positive association between proto-musical calling and swinging is driven by two contrasts—that between gibbons and hominids and between apes and Old World monkeys (Fig 1: contrasts 106 and 90). This is understandable considering that there are nearly no other brachiating primates in the rest of the primate tree (Fig 1). Thus, although the significant positive association of swinging with musical calling observed here is contingent upon methodological assumptions, a more complete sampling of gibbon species will likely improve the resolution of this conditional association. Interestingly, the methodological discordance, that seems to only separately highlight these alternate forms of aerial locomotion, entirely disappears when the two mutually exclusive measures are simply added together (Fig 2).

Perhaps the most illustrative inverse-example to our origins scenario is the case of cheek-pouch monkeys (subfamily *Cercopithecinae*) few of which are musical, leapers, or monogamous (Rowe and Meyers, 2017). Evidently, in their transition to a increasingly terrestrial existence (esp. *Papionini*), they lost all three of these traits. Only their hominoid relatives retained these traits long enough to find new adaptive functions as manifested in the swinging facilitated frugivory of socially monogamous lesser apes. While it likely required millions of years to fully unravel, the relatively recent radiation of these cercopithecines seems to have largely eroded the interdependent suite of arboreal specializations characteristic of their anthropoid progenitors.

Although the relationships we uncovered are robust under a number of different model compositions, they are admittedly largely driven by relatively few data-points—fewer than ten percent of the data drive the positive correlations. Furthermore, these contrasts correspond to branching times (Springer *et al*., 2012) that average to well over ten million years old. It seems likely that this co-evolution is slow forming but could also decouple if one or the other trait was atrophied. Also, it seems that monogamy, shown to co-vary with ARDI previously (Schruth, Templeton and Holman, under review), could further play an interesting role as part of a three way co-evolution. Familial acquirement of such precarious locomotion strategies (e.g. group crossing of canopy-gaps) may not only forefend predation of kin, but could have so radicalized the evolution of arboreal ranging logistics, that *efficient signaling* of any congruent cognition might also have been incentivized.

This selective influence of precarious, time sensitive locomotion could apply to many other animals besides primates—songbirds, hummingbirds, cetaceans, bats, and arthropods all arguably could be considered to have proto-musical calls (McDermott, 2008; Hoeschele *et al*., 2015), and all of whom either fly or swim. While the more aerial and terrestrial varieties above tend to land on thin terminal branches and slender grasses, the precise location of the surface for deep water diving mammals could have similarly unknown or otherwise challenging landing parameters. This could be particularly true for whales who feed on phytoplankton bloom driven food webs near polar ice sheets but must sometimes breath using polynyas. While it is known that species occupying habitats such as forest canopy or ocean depths use acoustic communication to efficiently overcome visual and olfactory obstructions (Slater, 2000), other forces are also likely at work as the calls of the orders listed above tend to go beyond just conveying location and identity. Mating (Darwin, 1871) and dominance (Hoeschele *et al*., 2010), perhaps in combination, could have selected for even more complex and elaborate calling patterns. As mentioned, we believe that the uncertainty of secure landing conditions alone could have provided substantial selective pressures for the co-evolution at these protracted evolutionary rates.

As we have shown, in non-hominin primates, arboreality, and thereby locomotion, appears to relate to musical calling. This pattern becomes complicated when considering our own genus which is much more terrestrial and musical than our semi-arboreal and less musical hominoid relatives (gorillas, chimps, and orangutans). That is, our parallel proposal that a more human-like musicality accompanied the hominin shift to terrestriality runs counter to the trend of the rest of the primate order. How is it that three other genera of hominoid failed to inherit the likely arboreal and musical mating system that the hylobatids seemed to have retained through the Miocene? The relatively recent discovery of *Ardipithecus ramidus*, a putative singer (Clark and Henneberg, 2017), indicates that arboreal locomotion, in the form of above-branch palmigrade clambering, may have been practiced as recently as four million years ago (Lovejoy, 2009; Lovejoy *et al*., 2009). Indeed, it is possible that this species (and presumably other Australopithecines) may have even slept in trees up until only a couple of million years ago (Fruth, Tagg and Stewart, 2018). It also appears terrestriality was something that evolved in parallel in multiple hominids (Larson, 1998; Lovejoy, 2009). Gorillas and chimps for example both became much more terrestrial and independently began knuckle walking millions of years after their divergence, perhaps due to increasingly dry conditions across the sub-continent (deMenocal, 2004).

So if an increase in terrestriality, and corresponding decrease in arboreality, primarily drives the *loss* of proto-musical calling, what is it about *Homo* that instead *promoted* musical behavior? It is possible that ballistics provides the answer. Accurate throwing (e.g. rocks, spears), the temporal reverse of catching (e.g. terminal branches) could pose similar selection pressures to aerial locomotion such as suspensory armswinging (Schruth, 2006). Humans throw things from great distance, with high momentum, and more accurately than any other species (Bingham, 1999). More generally however, tool use *is* known to be one of the primary defining characteristics of the genus Homo. The main evidence, dating back to Middle Paleolithic, abounds in the form of stone tool industries (Semaw *et al*., 1997), which could have co-opted the Miocene adaptations of suspensory arm-swinging for associated precision hammering. Wooden spears, unlikely to preserve for many thousands of years, nevertheless show up at least more recently (Thieme, 1997). Thus, even if we are not certain about brachiation driving musical calling in hominoids, it is possible that precision arm swinging, or more fine-motor skills for tool-making, engendered a suite of neurological changes that overlapped with an increasingly complex musical calling. Hominin dominance over seasonal resources (e.g. herds of game) could be derived from analogous behaviors of hominoids (e.g. over fruiting terminal branches) tens of millions of years previously—and both may have acted as evolutionary inducers of salient acoustic displays sharply directed (Searcy and Beecher, 2009) towards conspecific resource competitors.

Singing requires micro-athletic mastery over fine muscles (Nettl, 1983; Sacks, 2007) in the vocal apparatus as well as memory to match previous acoustic gestures with current utterances and to plan future such gestures, as has been suggested previously (Roederer, 1984). Subconscious pattern matching between disparate orbital inputs could modulate rectus muscle control of eye position in the ocular cavity thereby actuating stereoscopic vision for late-locomotor-bout, and potentially high-speed, substrate encounters. Aside from hand-eye coordination (e.g. ocular muscle coordination with distal-limb grasp-placement adjustments), another possibility includes breathing control (Hewitt, MacLarnon and Jones, 2002). Further extrapolations of musical behavior serving as a (non-vision based) motor control signal include that for the fine muscles of the fingers perhaps for intricate tool making by hominins. It is further tempting to speculate that performance drumming aspects of *rhythmic* musicality could signal related precision butchering abilities (Jordania, 2008) to other long-distance scavenging parties of hominins dispersed across these more open and arboreally sparse settings.

Humans, by themselves, constitute nearly the entirety of the terrestrially musical creatures on earth, making a solution to the evolutionary puzzle so challenging—we represent only a minority, an extreme outlier datum, among thousands of mostly non-terrestrial examples. There have been interesting explorations of understanding human music as derivative of more recent human adaptations such as rhythmic locomotion (Larsson, Richter and Ravignani, 2019) across earth’s two-dimensional surface (Mithen, 2006) or in association with later-developing faculties such as language (Livingstone, 1973; Pinker, 1997) or dance (Hagen and Bryant, 2003). While a counter-argument regarding the possible confounding with language origins could be made, our built-in requirement for redundancy (in ARDI) makes scenarios invoking co-evolution with the far less repetitive, referentially linguistic forms of communication less compelling. Our results instead ought to inspire consideration of the tens of millions of preceding years of three-dimensional arboreality in anthropoids, suspensory armswinging in hominoids, and ballistics of hominins all of which likely eventually enabled re-terestrialzation (Ishida, 2006) and hunting of associated game (Calvin, 1983). A proposed transition from precision limb landing, on tenuous branches, followed by precision hammering upon thin blade faces, for forging tools, is strongly evidenced by the near-unanimous arboreal affinities of extinct and extant primates and the scores of archaeological sites documenting hominin lithic productivity. This historical sequence fortifies a continuous adaptive co-evolutionary scenario from the Paleocene to the late Pleistocene.

In sum, our findings regarding the potentially three-way coevolution between locomotion, social monogamy, and our melodically-cognisant proto-musicality metric suggest that the curious case of human music has deep primate roots (Schruth, 2020). These roots plausibly derive from ancient patterns of subsistence based in precarious parabolic leaps, swings, and ballistic arches—all of which require last-minute fine-tuning adjustments in the wrist and fingers as well as high levels of coordination with the small muscles of the eye. Finally, if this arboreal, branch-dominance based locomotion evolved with more melodic calling (Schruth, Templeton and Holman, under review), then a shift to terrestrial size-dominance may have instead engendered more deep-toned and perhaps group-conducive, rhythmic musciality (Merker, 1999). This two part evolution of more delicate melodic aspects first, followed by more rugged rhythmic aspects second, corresponding to our hominoid to hominin transition between two drastically different habitats, may help to better illuminate the enduring enigma and astonishing uniqueness of human music.

